# Origin and evolutionary dynamics of multi-drug resistant and highly virulent community-associated methicillin-resistant *Staphylococcus aureus* ST772-SCC*mec* V lineage

**DOI:** 10.1101/2020.08.24.265801

**Authors:** Yamuna Devi Bakthavatchalam, Karthick Vasudevan, ShomaVinay Rao, Santosh Varughese, Priscilla Rupali, Maki Gina, Marcus Zervos, John Victor Peter, Balaji Veeraraghavan

## Abstract

**Background:** Community-associated methicillin-resistant *Staphylococcus aureus* (CA-MRSA) are increasing in prevalence across the world. However, studies on the molecular epidemiology and the genomic investigation of MRSA are limited in India.

**Objectives:** To understand the molecular epidemiology of MRSA and to reconstruct the origin and evolution of *S. aureus* belonged to the sequence type (ST772).

**Methods:** A total of 233 non-repetitive MRSA isolates were screened for the presence staphylococcal cassette chromosome (SCC*mec*) types, multi-locus sequence types (MLST) and staphylococcal protein A (*spa*) types. Whole genome sequence data of ST772-SCC*mec* V (n=32) isolates were generated and analysed along with the publically available ST772-SCC*mec* V (n=273) genome.

**Results:** ST772 (27%), ST22 (19%) and ST239 (16%) were found as the predominant STs. Analysis of the core SNPs using Bayesian time scaled phylogenetic analysis showed ST772-SCC*mec* V was emerged on the Indian subcontinent in 1960s. The acquisition of integrated resistance plasmid (IRP) in the ST772-SCC*mec* V lineage during 1990s, fixation of SCC*mec* V (5C2) and the double serine mutations (S84L, S80Y) appears to have played a key role in the successful expansion. The IRP carries the loci for multiple antibiotic resistant genes: beta-lactam (*bla*Z), aminoglycosides (*aph*A3-*sat*-*aad*E), macrolide (*mphC*), macrolide-lincosamide-streptogramin B (*msrA*) and bacitracin (*bac*A, *bac*B).

**Conclusion:** The Panton Valentine Leukocidin (PVL) positive ST772 and ST22 MRSA lineages are observed in the hospital settings. ST772-SCC*mec* V has the multi-drug resistance trait of hospital-associated (HA) MRSA and the epidemiological characteristics of CA-MRSA. The antimicrobial use pattern may have driven the spread and survival of ST772 MRSA in hospitals.

## Introduction

Methicillin-resistant *Staphylococcus aureus* (MRSA) is of significant clinical concern in causing both community-associated (CA) and healthcare-associated (HA) infections. MRSA was first described in 1996.^1^ It has merged through the acquisition of different staphylococcal cassette chromosome *mec* (SCC *mec*) carrying methicillin resistance determinant *mec*A.^2^ The HA-MRSA strains carries the SCC*mec* types I, II or III and are often multi-drug resistant. Typically, the CA-MRSA strains carries the SCC*mec* types IV or V, a Panton-Valentine leukocidin (PVL) toxin and are frequently susceptible to non-beta-lactam antibiotics.^3^

Most of the HA-MRSA clones belong to the clonal complex (CC), CC5, CC8, CC22, CC30 and CC45.^4^ In contrast, CA-MRSA clones are geographically more diverse. In the Asia-pacific region, five distinct sequence types (STs); ST59, ST30, ST72, ST8, and ST772 are the predominant CA-MRSA clones and have often spread to hospitals.^5^ An increasing resistance to non-beta-lactam antibiotics have been reported in CA-MRSA.^6^ In India, MRSA accounts for about 37% of *S. aureus* infections.^7,8^ However, there are only limited studies on the molecular epidemiology of *S. aureus*.

In early 2000, most of the Indian MRSA isolates belonged to the ST239-SCC*mec* III lineage and were restricted to hospitals. ^9-12^ In 2004, a novel CA *S. aureus* clone, ST772-Methicillin-susceptible *S. aureus* (MSSA) isolated from hospitals in Bangladesh and ST772-SCC*mec* V MRSA (Bengal bay clone) isolated from the community and hospitals in India has been described.^12,13^ During the same year, a variant of epidemic MRSA (EMRSA-15) carries PVL, SCC*mec* IV and belongs to ST22 has also been reported in India.^12^ After 2004, ST772 and ST22 has increased in prevalence to become the dominant STs in both community and hospital infections in India.^14-19^ In the last decade, ST772-SCC*mec* V has proven to be successful at disseminating into hospitals, begin with localized expansion and rapidly progressing to an intercontinental spread.^20,21^

Genomic studies have brought both understanding about the evolution of *S. aureus* and new insights into the genetic determinants contributing to its spread and survival. The well-studied HA-MRSA genomes are mostly restricted to the Hungarian clone (ST239-SCC*mec* III) and EMRSA-15 (ST22-SCC*mec* IV).^22,23^ Other well-studied CA-MRSA genomes are the USA300 clone (ST8-SCC*mec* IV) and a live-stock associated MRSA, the ST398-SCC*mec* V lineage. ^24, 25^

The present study objectives were i) to provide preliminary data for isolate selection that belongs to ST772-SCC*mec* V lineage for further genomic investigation, 233 MRSA isolates were screened for genotypes using SCC*mec* typing, multi-locus sequence typing (MLST) and staphylococcal protein A (*spa*) typing, ii) to reconstruct the origin of ST772-SCC*mec* V isolates collected between 2004 and 2018, iii) to establish a high resolution phylogeny and evolutionary analysis of this emerging lineage, iv) to trace the genetic changes associated with its evolution and success and v) to characterise the pangenome of the ST772-SCC*mec* V lineage.

## Materials and methods

### Bacterial isolates

Non-repetitive MRSA (n=233) isolates (one isolate per patient) recovered from the blood culture samples during 2013-2018 were included in this study. MRSA isolates were identified using standard bacteriological methods including gram staining, culture and tube coagulase test.^26^ The study was conducted in a 2600 bedded tertiary care, Christian Medical College and hospital, Vellore, India and approved by the Institutional Review Board of Christian Medical College, Vellore (IRB.Min.no. 10643, dated on April, 2017).

### Molecular characterisation of MRSA

SCC*mec* types were determined using multiplex PCR as described by Zhang *et al*.^27^ The subtypes of SCC*mec* IV were identified as described previously.^28^ MLST of *S. aureus* was performed as defined by Enright *et al*.^29^ STs and CC were assigned according to the MLST database (https://pubmlst.org/saureus/). *spa* typing was performed as described by Harmsen *et al*.^30^ Assignment of the *spa* types were carried out using the public *spa* type database Ridom SpaServer (www.spaserver.ridom.de).

### Genome sequencing and assembly

A representative subset of 32 isolates of the predominant ST772-SCC*mec* V lineage were further analysed using whole genome sequencing and compared with the available published genomes. Then, the genetic lineage, virulence and resistant determinants were described.

#### Ion Torrent sequencing

Whole genome sequencing was performed using IonTorrent PGM platform (Life Technologies, Carlsbad, CA). The sequencing library was prepared using 200ng of genomic DNA which was sheared to a fragment length of 400bp. Library preparation and sequencing were performed according to the manufacturer’s instruction using Ion Plus Fragment Library Kit (Life Technologies, California, United States). De novo assembly of the raw reads generated from ion torrent were assembled using AssemblerSPAdes software v4.4.0.1 in Torrent suite server version 4.4.3.

#### Genome annotation and downstream analysis

The genome was annotated using the RAST server (Rapid Annotation using Subsystem Technology, http://rast.nmpdr.org)^31^ available from the PATRIC database (Pathosystems Resource Integration Centre, https://www.patricbrc.org)^32^ and the National Center for Biotechnology Information (NCBI) Prokaryotic Genome Automatic Annotation Pipeline (www.ncbi.nlm.nih.gov/genomes/static/Pipeline.html). SCC*mec* types were determined using the SCC*mec* finder database.^33^ MLST of the isolates were reconfirmed using the MLST finder (https://cge.cbs.dtu.dk/services/MLST/) which is available from the Centre for Genomic Epidemiology database.

#### Genome analysis for virulence and resistant genes

Virulence factors were identified using virulence factor database (VFDB)^34^ and the acquired virulence genes using VirulenceFinder (https://cge.cbs.dtu.dk/services/VirulenceFinder/). Acquired antimicrobial resistant (AMR) genes were identified by using the following different AMR databases; BLAST search against comprehensive antibiotic resistance database (CARD)^5^, ResFinder 3.2 (https://cge.cbs.dtu.dk/services/ResFinder/), KmerResistance 3.1 (https://cge.cbs.dtu.dk/services/KmerFinder/), Mykrobe predictor^36^ and Pathogenwatch (https://www.sanger.ac.uk/science/tools/pathogenwatch). Insertion of the phage genome was identified using PHASTER.^37^ Pathogenicity Islands were identified using the Pathogenicity island database. ^38^

#### Genome comparison

The original ST772-SCC*mec* V global data of Steinig *et al* ^39^, Prabhakara *et al* ^40^ and Balakuntla *et al*^41^(Supplementary Table 1) were supplemented with the genome of an additional 32 isolates sequenced in this study collected during 2013-2018. Sequence reads of *S. aureus* ST772 isolates (n=273) were retrieved from the NCBI and European Nucleotide Archive (ENA) database and assembled with SPAdes v.3.12^42^. DAR4145 (accession no. CP010526) deposited from Mumbai was used as the reference genome.

The capsular biosynthesis gene (*cap*A-G and *cap*L-P), an accessory gene regulator (*agr*BDCA), a DNA repair protein (*rad*C), the type-I restriction-modification system (*hsd*R) and the type III restriction system were analysed for mutations. The effect of amino acid substitution on the protein function was predicted using the sorting intolerant from tolerant (SIFT) score.^43^

### Single nucleotide polymorphism (SNP) based phylogenetic analysis

Phylogenetic analysis was performed against the publically available global genomes. The genomes were aligned to the reference genome using BWA mem algorithm.^44^ Variants were called using Snippy v.4.5.1.^45^ and FreeBayes ^46^ with a minimum base quality of 20, a minimum read coverage of 10X and a 90% concordance at a locus for a variant were selected. A whole SNP genome alignment and core SNPs genome alignment of all genomes was generated with snippy-core. Core SNPs were defined as being present in all isolates and were extracted using snippy-core with default settings was used to infer a phylogeny. The recombination free core genome SNP alignment file generated by Gubbins was used as the input to compute the mean evolutionary rate of the genomes and time of the most recent common ancestor (MRCA).^47^ The RaXML-NG 0.5.0 was used to generate a maximum-likelihood (ML) tree based on 6344 variant sites in the core genome. ^48^ The general time reversible model (GTR + Γ) of nucleotide substitution was implemented with ascertainment bias correction (Lewis) and 100 bootstrap replicates as suggested by Steinig *et al*.^39^ The tree with the highest likelihood was midpoint rooted and visualised with iTOL.^49^ The study isolates and the clades were assigned to previously described lineages.^39^ A hierarchical Bayesian analysis of population structure (hierBAPS) clustering model was used to support phylogenetic groupings by using iterative clustering to the depth of 10 levels and a pre-specified maximum of 20 clusters. ^50,51^

### Bayesian time scaled phylogenetic analysis

Temporal signal in the ML phylogeny for ST772 *S. aureus* was investigated using TempEST v 1.5.1 (http://tree.bio.ed.ac.uk). The relationship between root to tip distance and the time of isolation were analysed. Root-to-tip regressions (R2) may indicate the evolutionary rate and probable outliers among the isolates. The temporal phylogenetic structure was determined using Bayesian Evolutionary Analysis by Sampling Trees (BEAST) v1.10.4 package after the initial assessment of the temporal signal.^52^ Sources and date of isolation of 305 ST772 *S. aureus* isolates used for temporal and BEAST analysis are listed in Supplementary Table 5. Evolutionary rates and tree topologies were determined with the generalized time reversible (GTR) substitution models with gamma distributed among-site rate variation with four rate categories (GTR+Γ4+I). The strict (assumes that branches in the tree evolves at constant evolution rate) and uncorrelated relaxed molecular clocks (assumes that each branch in the tree have independent evolution rate) were implemented with three demographic models, Constant size, Exponential and Bayesian skyline coalescent tree priors. The MCMC chain was run for 500 million steps with sampling of 20000 generations. We performed an additional duplicate run of 500 million steps and combined the log using LogCombiner v1. 8.2 when the effective sample size (ESS) estimates did not reach 200.^53^ The convergence and mixing of each run was manually inspected Tracer.v.1.7 to ensure that all the runs converged to an ESS of >200.^54^ To determine the best-fitting coalescent model to describe changes in effective population size over time, log marginal likelihoods were calculated using path sampling and stepping stone sampling methods. Finally, Bayes factor was used to determine the best fit model^55^, a burn in of 20% was discarded from the runs and the maximum clade credibility (MCC) tree was generated using Treeannotator v.1.8.2.^56^ BEAST output was analyzed using Tracer v1.7, with uncertainty in parameter estimates reflected as the 95% highest probability density (HPD). The annotated phylogenetic tree was visualized using FigTree v.1.4.6 and the Bayesian skyline plot was reconstructed using Tracer.v.1.7.^57,58^

### Pangenome analysis

Pangenome analysis of ST772 *S. aureus* (n=305) was performed using Roary v. 3.11.2 with default settings.^59^ The genomes were annotated using Prokka v.2.1.4.^60^ The genes that were common in all compared strains (core genes) and accessory genes were extracted and was used to contruct a phylogenetic tree. Core genes (99% ≤ strains ≤ 100%), soft core genes (95% ≤ strains < 99%), shell genes (15% ≤ strains < 95%), cloud genes (0% ≤ strains < 15%) and total genes (0% ≤ strains ≤ 100%) were calculated. In addition, the presence or absence matrix representing accessory genes were graphed and visualized using roary_plots.py python script (https://github.com/sangerpathogens/Roary). Mobile genetic elements present in the genome were identified using comparative genomic analysis and manual curation. Accessory genome analysis was divided into *S. aureus* Pathogenicity Island (SaPI), prophage, plasmids, SCC*mec* elements and transposons that are known to contribute major source of variation between *S. aureus* genomes.^22^ Mobile genetic elements (MGEs) in the genomic sequences were analysed using comparative genomic analysis and manual curation. IRP, transposons and insertion sequence (IS) of DAR4145 were used as the reference sequences. In addition, IS mapper and was used for the screening of IS elements. Detailed comparisons of the genome sequences were performed on the de novo assemblies using BLASTN. Hypothetical proteins are those did not have any matches either with core or MGEs of reference genome sequence (DAR4145). The core genome elements were determined using Spine V0.3.1 with default cutoff (Minimum nucletotide percent identity of 85% and 100% minimum threshold to identify as core).^61^ Additionally, accessory genomic elements were identified using AGEnt v0.3.1 that were not identified as core genomes by Spine.^61^ Further, the accessory genomes from all the isolates were combined and functional annotation was determined using eggNOG-mapper.^62^

## Results

### Preliminary molecular characterisation of MRSA isolates

SCC*mec* V was the predominant gene in 42% (n=98) of the isolates followed by SCC*mec* III in 27% (n=63), SCC*mec* IV in 13% (n=31), SCC*mec* I in 7% (n=16) and SCC*mec* II in 3% (n=7) of the isolates. The SCC*mec* IV subtypes were IVa (n=9), IVc (n=7), IVd (n=14), and IVh (n=1). Notably, eighteen isolates had multiple SCC*mec* types; fifteen isolates carried SCC*mec* III and SCC*mec* V and one isolate was identified with each of the following SCC*mec* I and SCC*mec* IVd; SCC*mec* IVd and SCC*mec* V; SCC*mec* IVa and SCC*mec* V.

MLST analysis revealed ten different CC and three singletons (ST616, ST1598, ST1947) (Fig 1). Among the CCs, five high-risk clones (HRC) including CC5, CC8, CC22, CC30, and CC45 were observed. Three major CCs; CC1 (ST1, ST772), CC22 (ST22, ST217, ST636, ST896, ST1037, ST2371, ST3976) and CC8 (ST8, ST239, ST368, ST630, ST1803, ST3324) accounted for 82% (n=192) of the isolates. Other minor CCs and their associated STs were given in Table S1. A total of 32 different STs were identified. The most common were ST772 in 27% (n=63), ST22 in 19% (n=44), ST239 in 16% (n=37) and ST368 in 9% (n=20) of the isolates (Fig 1). Although, the STs were observed to be diverse, most of them were recognised as a SLV or DLV of the dominant STs (Table S2). Thirty distinct *spa* types were identified. The *spa* type t657 was identified in 34% (n=73) of isolates followed by t425 in 18% (n=34), t037 in 12% (n=25), t030 in 8% (n=18) and t852 in 10% (n=17). The remaining 25 different *spa* types were found in fewer than five isolates (Fig 1, Table S3).

**Fig 1:**
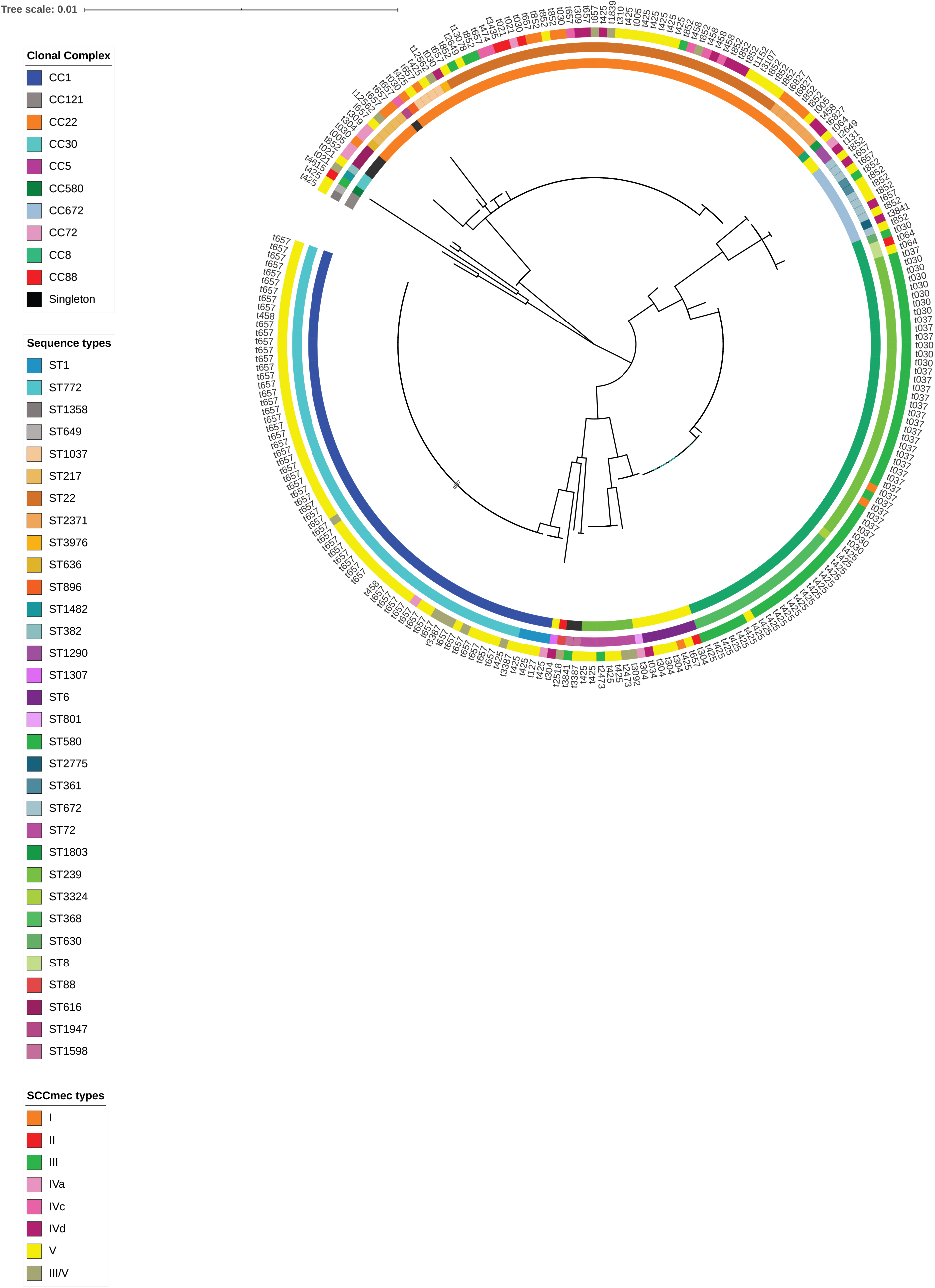
a) Maximum likelihood phylogeny was constructed based on the seven house-keeping genes of *S. aureus*. The tree was marked with the meta-data using interactive tree of life (iTOL.v.4). The inner most ring represents clonal complexes (CCs), the second ring denotes sequence types (STs), the third ring represents SCC*mec* types and the outer most ring represents the *spa* types. CCs, STs and SCCmec types are colour coded as mentioned in legends.

ST772-SCC*mec* V-t657 lineage was observed in 22% (n=52), ST239-SCC*mec* III lineage sharing the *spa* types t037 and t030 in 15% (n=35) and ST368-SCC*mec* III-t425 lineage in 8% (n=19) of the isolates were identified. In contrast, SCC*mec* and *spa* types were highly diverse among the ST22 MRSA strains (Fig 1, Table S3).

### Genomic analysis of *S. aureus* ST772-SCC*mec* V

#### Phylogenetic analysis

A representative subset of 32 ST772-SCC*mec* V MRSA isolates characterized in the preliminary screening were included and compared with the genomes that are available in the public database. The country and the year of isolation of ST772 *S. aureus* isolates are given in the Supplementary Fig 1. ST772-SCC*mec* V genome data was comprised of MRSA (n=270) and MSSA (n=34) isolates. Ten basal strains (4 MSSA, 6 MRSA) were also included. Mapping of ST772 S. aureus isolate against the reference genome resulted in an alignment of of 6344 core SNPs, which were used for phylogenetic inference. A maximum likelihood phylogenetic tree generated from the core genome SNP is presented in Fig 2a. ST772-SCC*mec* V isolates were grouped into two distinct clades; clade I represented basal strains, the earliest was to found to be MSSA which was isolated from India and Bangladesh in 2004. The clade II was recognised as a globally distributed monophyletic group and diverged into three different subclades (A, B, C). Majority of the isolates clustered in clade IIB (56%, n=169) which were considerably more diverse and dominant than early branching clade IIA (24%, n=73) and emerging clade IIC (17%, n=52) isolates. The phylogenetic comparison of Indian ST772 *S. aureus* isolates against isolates available from other countries showed that Indian isolates were distributed across all the subclades and also the clade containing basal strains.

**Fig 2:**
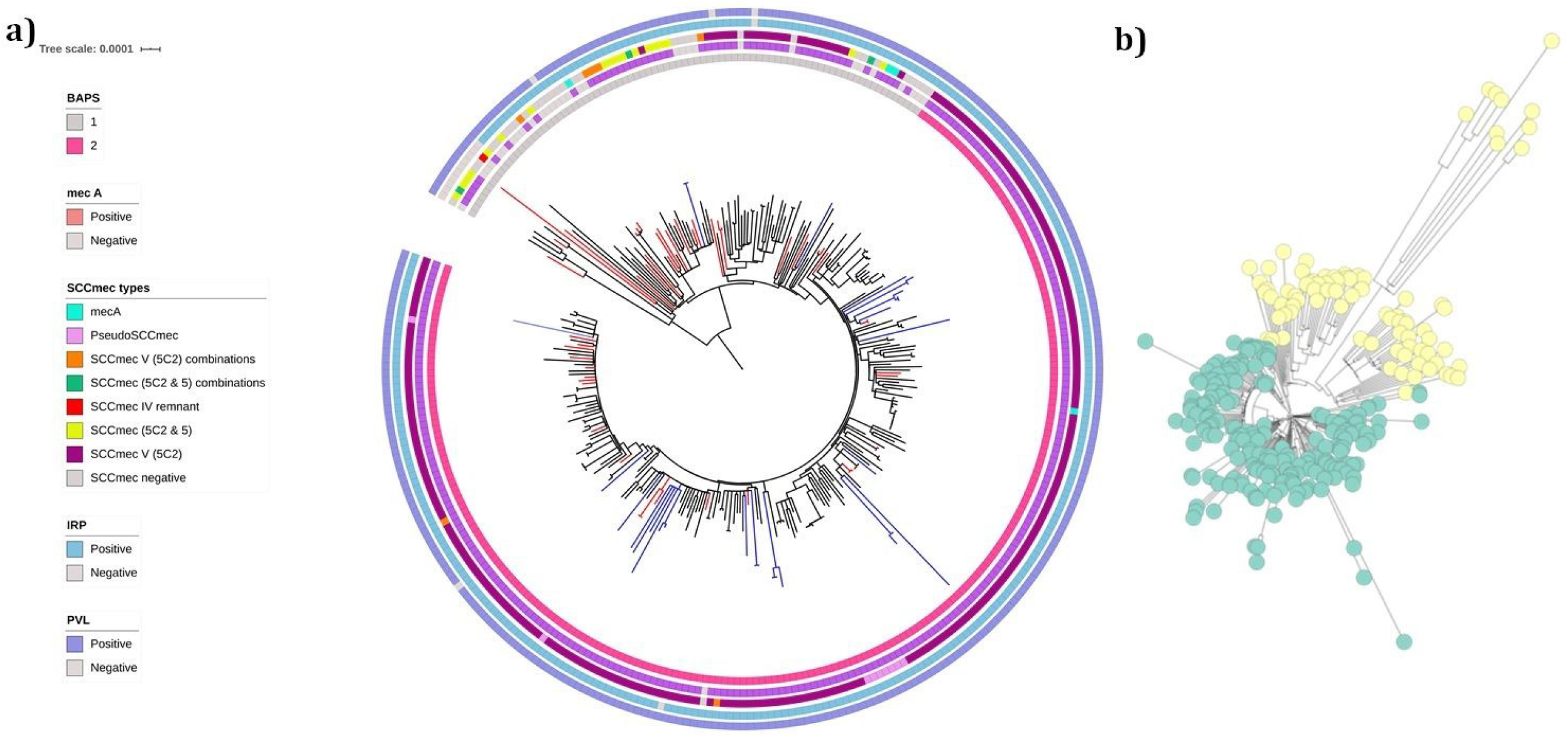
a) Maximum likelihood phylogenetic tree was constructed based on the core genome SNP of ST772 *S. aureus* lineage (n=305). The innermost ring represents BAPS clusters. The second, third, fourth and fifth rings denotes the presence/absence of *mec*A gene, presence of various SCC*mec* types, presence/absence of integrated resistance plasmid (IRP) and the PVL gene respectively. Isolates in the clades are colour coded; study isolates are in blue color and other Indian isolates are in red color. b) Bayesian Analysis of Population Structure (BAPS) analysis of ST772 *S. aureus* isolates showed the presence of two distinct sequence clusters (SC) based on the core SNPs. The basal strains and isolates of subclade IIC (yellow) are classified in SC1, while isolates of subclade IIb(green) are represented in SC2.

Population structure analysis using RhierBAPS revealed two distinct BAPS groups (Fig 2b). The basal strains and isolates of subclade IIA were classified as BAPS population cluster 1, while isolates of subclade II B and C were categorised as BAPS population cluster 2. In BAPS population cluster 1, sequence comparison of 83 isolates (basal strains and subclade IIA) against the reference genome DAR4145 identified 3306 core SNPs (among a core genome of 2.8 Mbp). Similarly, in BAPS population cluster 2, a total of 3678 core SNPs (among a core genome of 2.8 Mbp) were identified by aligning the sequences of 221 isolates (subclade IIB and C) against the reference genome.

#### Bayesian time scaled phylogenetic analysis

As there was a strong temporal signature for the observed mutations in the global ST772 *S. aureus* population, we explored the temporal structure of ST772 S. aureus isolates using Bayesian phylogenetic methods. The root to tip analysis revealed a strong correlation (correlation coefficient 0.6862 and R squared 0.4709) between the time of isolation and distance from root suggesting the temporal clock like evolution in the lineage (Fig 3). A maximum clade credibility tree was reconstructed based on the concatenated core SNPs and is shown in Fig 4. The mean substitution rate of the ST772 *S. aureus* population was estimated to be 1.16 ×10^−6^ substitution per site per year. The time-scaled phylogenetic reconstruction suggest that the MRCA of ST772 *S. aureus* with initial divergence was estimated to be from the year 1959 (95% HPD, 1936-1979). This was followed by the emergence of dominant clade II isolates and its subgroups in 1987 (95% HPD, 1981-1995). Bayesian hierarchical clustering using core SNPs segregated *S. aureus* ST772 isolates that had IRP and a variant of SCCmec V (5C2) into separate subclades (Fig 4). This infers that the ST772 *S. aureus* resistant subgroups expanded rapidly after 1987 and has since sustained.

**Fig 3:**
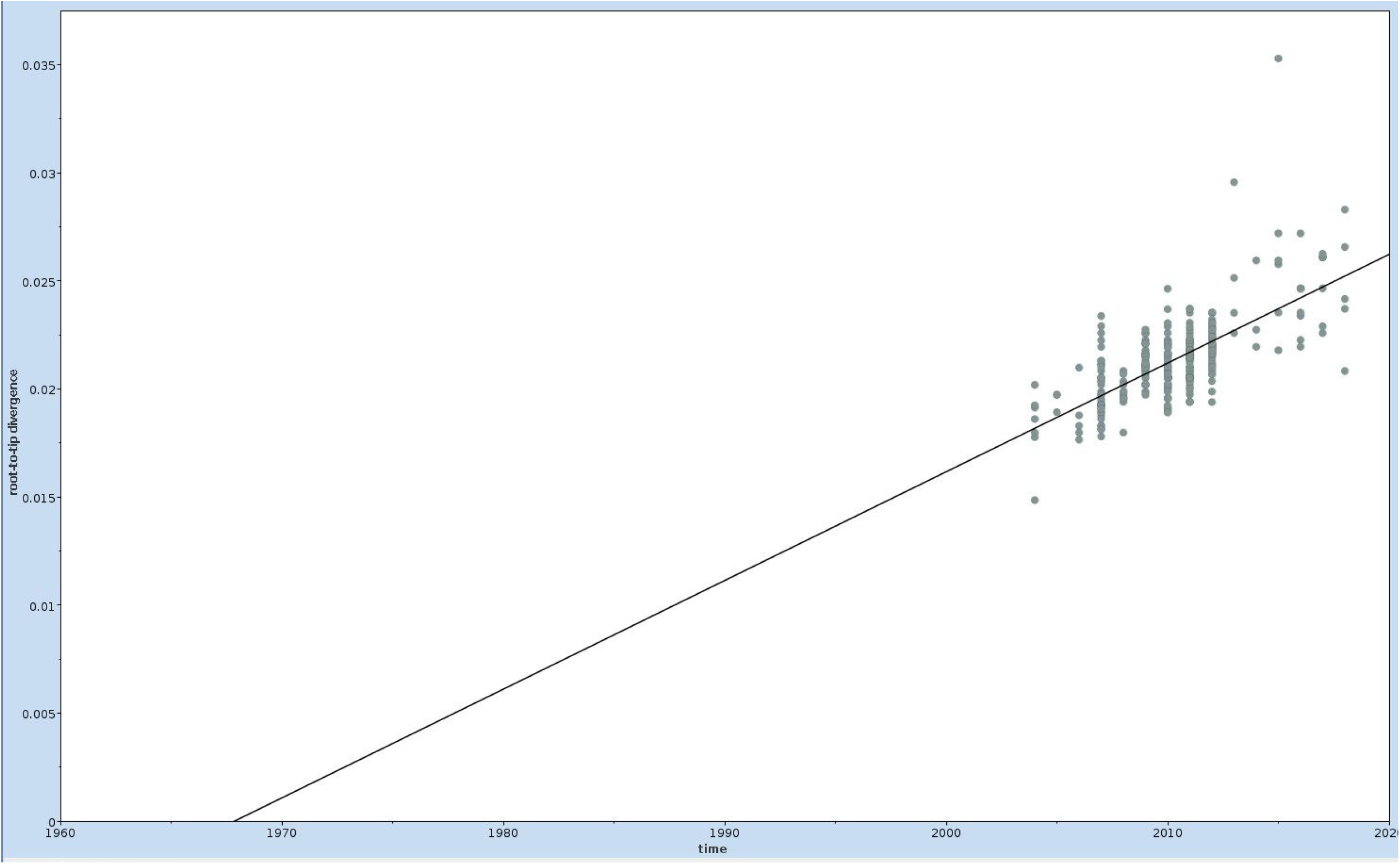
Root to tip genetic distance of the ST772 *S. aureus* isolates. The total number of SNPs present within each isolate compared to that in the index isolate was computed and plotted against the date on which each isolate was recovered. The graph includes regression lines with 95% CI for each group. Sizes of circles correspond to the number of isolates.

**Fig 4:**
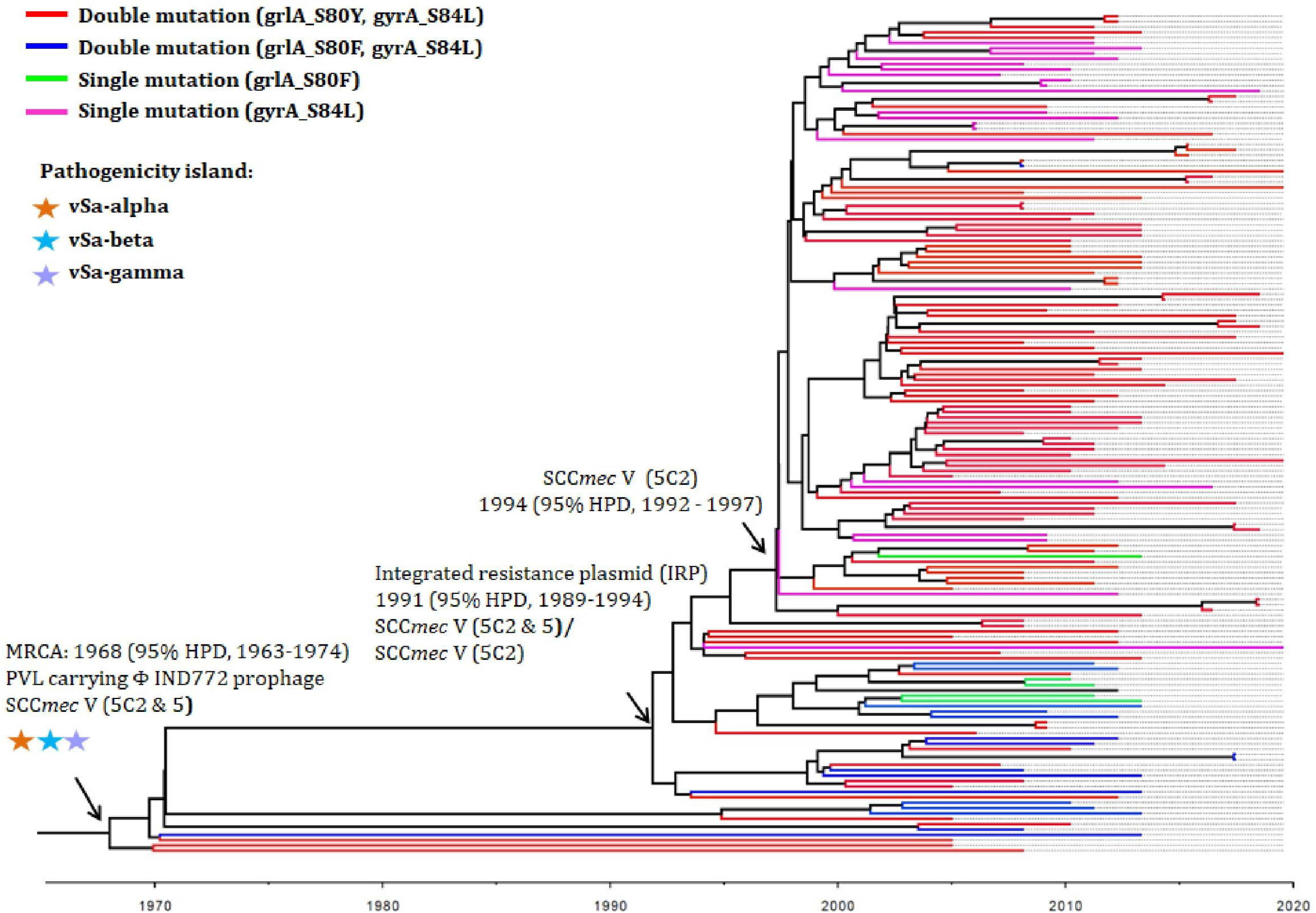
Evolutionary analysis of ST772 *S. aureus*. The maximum clade credibility tree was generated from the filtered single-nucleotide polymorphisms (SNPs) in the core genome of 305 *S. aureus* ST772 genome using BEAST analysis.

The generated BSP estimated the demographic changes of ST772 *S. aureus* genome (n=305) over time (Fig 5). The effective population size of ST772 was relatively stable from early the 1950s to the late 1970s. Notably, two mild expansion events were noticed. Initial event was associated with the acquisition of IRP, and SCCmec V (5C2) in 1987 followed by another event which replaced the double serine mutations *gyr*A (S84L) with *grl*A (S80F) by *gyr*A (S84L) with *grl*A (S80Y) in 1993. These genetic changes resulted in a sharp increase in ST772-SCC*mec* V effective population size in 1997. At this stage, the population size increased by one order of magnitude. After 2010, there was a sharp decline in the effective population size and recently there has been a noticeable plateau.

**Fig 5:**
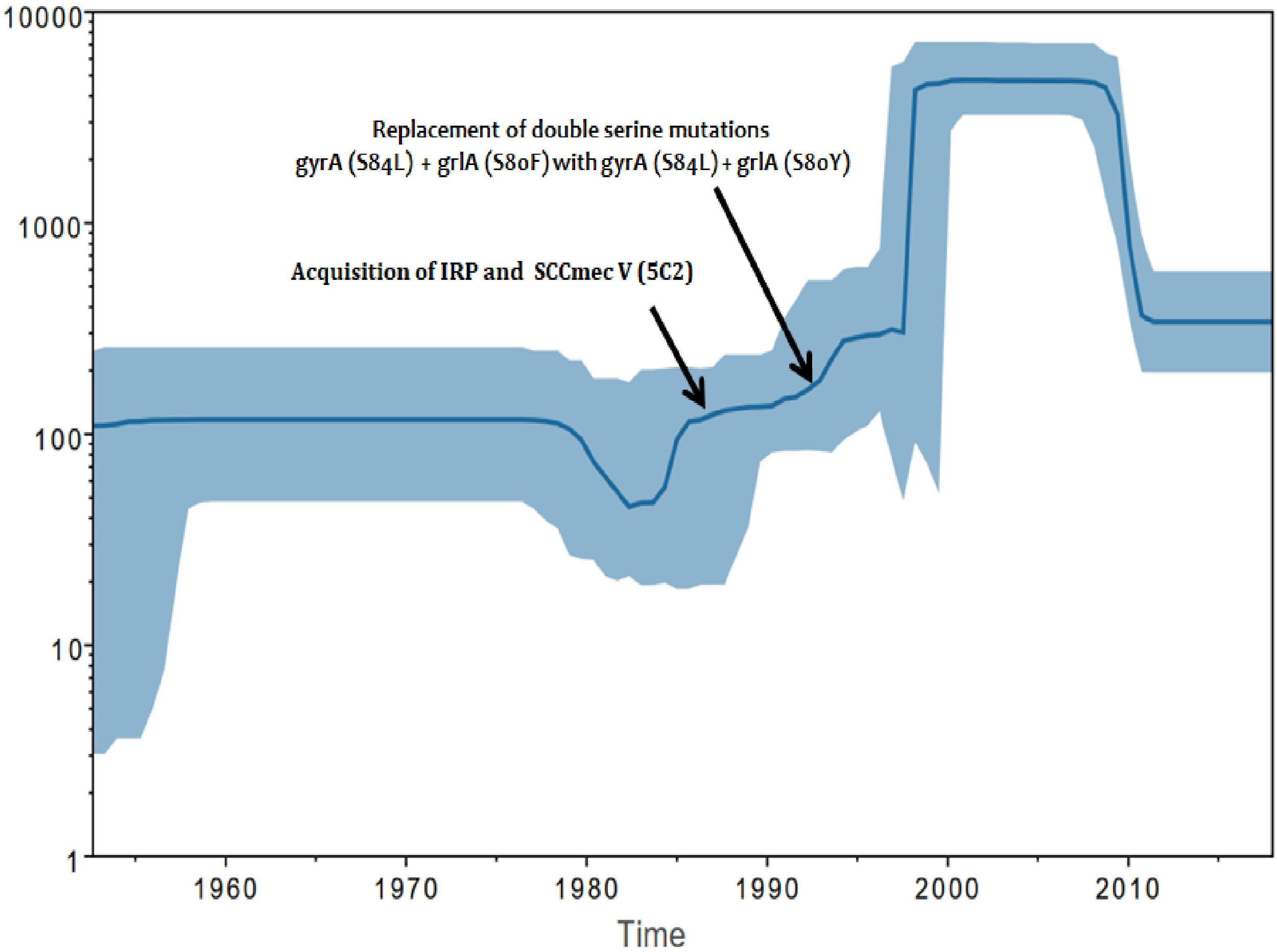
The Baesiyan skyline plot (BSP) analysis of ST772 *S. aureus* effective population size and it’s time for the most recent common ancestor. BSP was based on the strict clock’ coalescent framework analysis and was constructed using 305 sequences (representing all years and countries). X-axis represents time in years and Y-axis shows the effective population size. The blue thick line represents the median, while the blue band represents 95% highest posterior density (HPD) intervals.

#### Virulence factors

Virulence factors identified in ST772 *S. aureus* are given in the Table S3. Almost all of the isolates (99%, n=301) carried a 59.4 kb Φ IND772 prophage containing PVL-operon (*luk*S and *luk*F) and a staphylococcal enterotoxin A (*sea*). Three different genomic islands carrying various virulence factors were identified; a 41 kb, vSa-alpha genomic island containing an exotoxin cluster (*set*6, *set*7, *set*8, *set*10, *set*11, *set*12, *set*13, *set*14, *set*15); a 25.2 kb vSa-beta genomic island containing an enterotoxin gene cluster (*sec, seg, sei, sem, sen, seo*); and a 24.6 kb, vSa-gamma genomic island contains an extracellular fibrinogen binding protein (*ecb*), a fibrinogen binding protein (*fbp*), an alpha hemolysin (*hla*) and an exfoliative toxin A (*eta*) (Fig S2). All the three genomic islands were universally seen across the basal strains and isolates of globally disseminated clade II (subclades IIA, B and C) (Fig 6). In addition, all the isolates were found to carry *aur, scn, hld, ebs* and a biofilm encoding *ica* operon (*ica*ADBC) in the genome (Fig 6 and Table S3).

**Fig 6:**
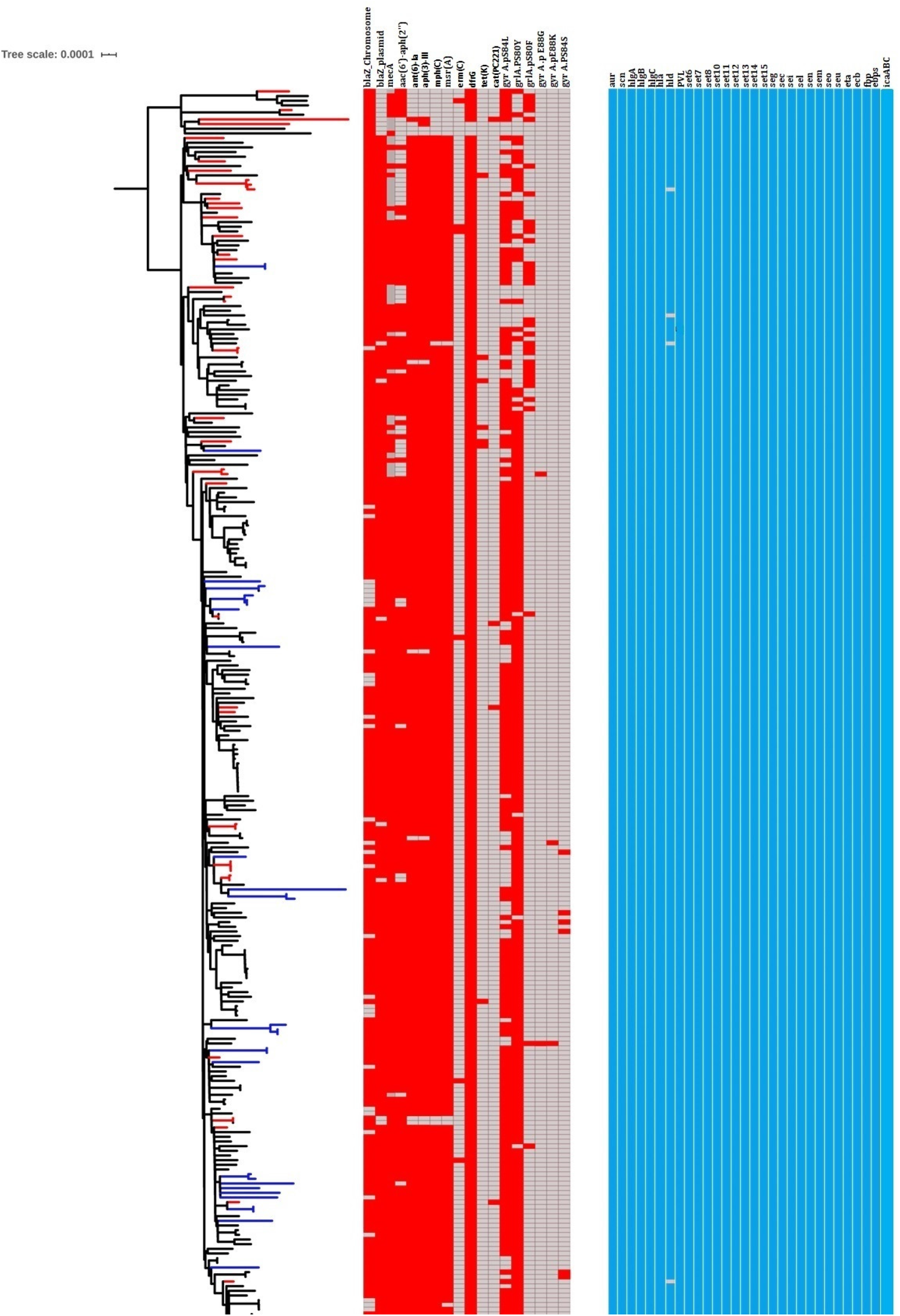
Distribution of antimicrobial resistant (AMR) genes and virulence genes identified among the clades of ST772 *S. aureus* isolates. The presence of acquired AMR genes as well as chromosomal mutations is denoted in red color. The presence of virulence genes are indicated in blue colour.

Gene sequences of *cap*A-G, *cap*L-P, *agr*BDCA, *radC*, type I (*hsd*R) and type III restriction system were highly conserved among the ST772 lineage. The substitutions and premature stop codons identified in these genes are listed in the Table S4. The substitutions in *cap*5B (K55N, K55Q) and *cap*5D(K7E) were predicted to affect the protein function with a SIFT score of 0.001.

#### SCC*mec* types

A 47.2 kb composite SCC*mec* V (5C2 and 5) and a 29.2kb SCC*mec* V (5C2) variant were recognised in 7% (n=21) and 76% (n=230) of isolates respectively (Fig 2a). All the basal MRSA (n=6) isolates harboured a large composite SCC*mec* V (5C2 and 5) element. The diversity of MRSA carrying SCC*mec* types decreased as ST772 clade II diverged into subgroups: clade IIA comprised of MSSA (n=28), SCC*mec* V (5C2 and 5, n=17) and SCC*mec* V (5C2, n=26); in contrast subclades IIB and IIC were exclusively carried SCC*mec* V (5C2) (Fig 2a). A transposon Tn4001 carrying aminoglycoside resistance (*aac*A-*aph*D) gene in SCC*mec* V was noted in 87% (n=264) of the isolates (Table S5). Various cassette chromosome recombinase genes (*ccr*AA, *ccr*AB2, *ccr*C), a high affinity potassium transporter (*kdp*), and a zinc resistance gene (czrC) were found to be integrated into SCC*mec* V (5C2 and 5) and SCC*mec* V (5C2) elements (Table S5).

#### Chromosomal mediated antimicrobial Resistance

Acquired genes and chromosomal mutations encoding for antimicrobial resistance is shown in Fig 6. Acquisition of Tn552 transposon-mediated beta-lactamase (*bla*Z) conferring penicillin resistance was identified in all of the isolates (Table S5). For the identified copies of *bla*Z, 6% (n=19) were carried on the chromosome, 25% (n=75) were carried on the IRP, and 69% (n=211) were carried in two location (one copy on the chromosome, and another on the IRP). Trimethoprim resistance determinants *dfr*G and *dfr*A were identified in the chromosomes of 99% (n=302) and 100% (n=305) of the isolates respectively. The most common mechanism of fluroquinolone resistance was the double serine mutations; *gyr*A (S84L) and *grl*A (S80Y, S80F). The mutations S84L with S80Y and S84 with S80F accounted for fluroquinolone resistance in 70% (n=215) and 9% (n=28) of the isolates respectively. The double serine mutation (S80F with S84L) was predicted to be replaced by S84L with S80Y in the early 1990s (Fig 4).

#### Acquired antimicrobial resistance

All ST772 isolates uniformly carried an IRP, except the basal strains (Fig 6). The IRP contained loci for multiple antibiotic resistant genes: beta-lactam (*bla*Z), aminoglycosides (*aph*A3-*sat*-*aad*E), macrolide (*mphC*), macrolide-lincosamide-streptogramin B (*msrA*) and bacitracin (*bac*A, *bac*B) (Fig S3). This genetic mobile element was recognised in 96% (n=292) of the isolates. Aminoglycoside resistant determinants (*aph*A3 and *aad*E) were identified in 95% (n=290) of the isolates (Table S5). The genes *mphC* and *msrA* were found in 96% (n=292) and 96% (n=291) of the isolates respectively (Table S5).

#### Pangenome analysis

The pangenome analysis of ST772 *S. aureus* genome identified 7428 orthologs with 2040 core genes (27.4%) and 5388 accessory genes (72.6%) (Fig 7). This included 283soft core genes, 376 shell genes and 4729 cloud genes. Among the accessory genes, 50% (n=2669) were hypothetical proteins. The isolate ERR217355 had high number of accessory genes (n= 2613), compared to others.

**Fig 7:**
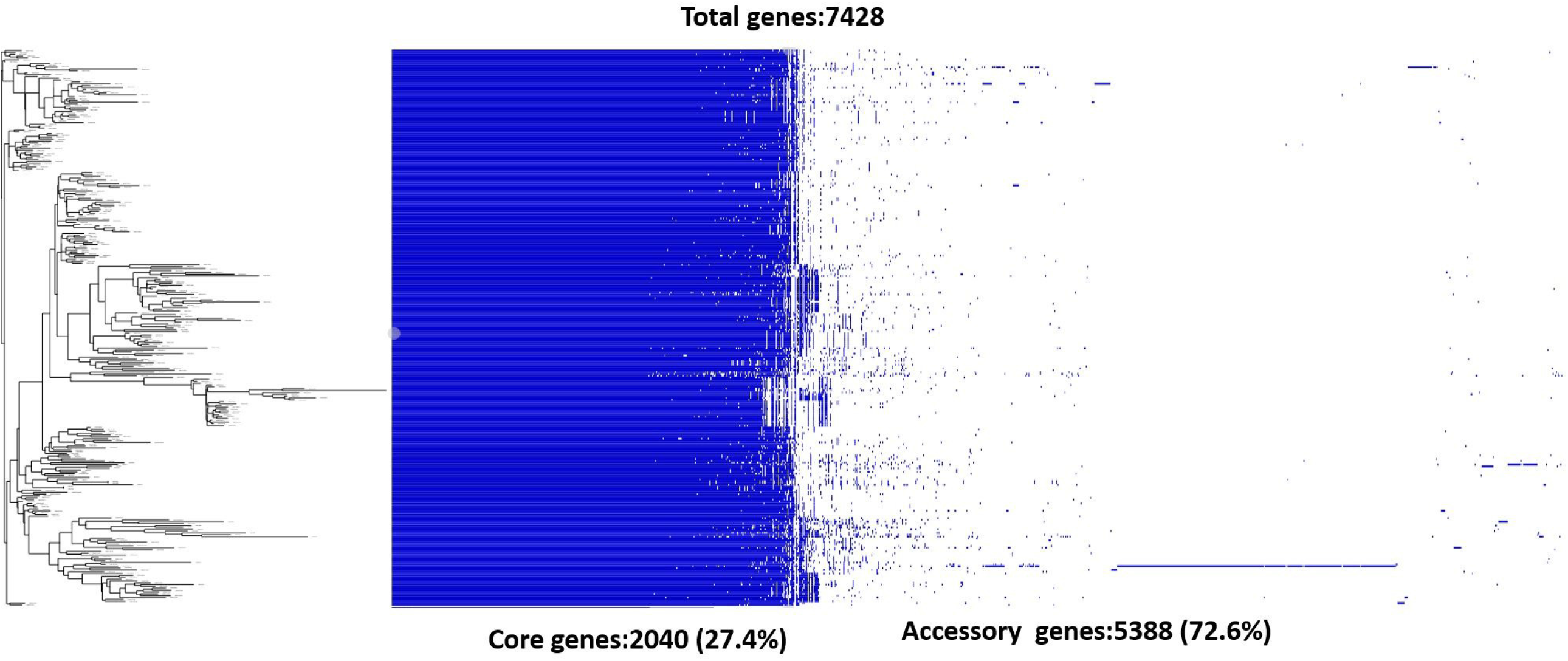
Pangenome analysis of ST772 *S. aureus* isolates using roary pipeline and visualised using Phandango, an interactive viewer for bacterial population genomics. Blue bar indicates the pan-genome of ST772 *S. aureus* isolates with 7428 annotated genes detected in all the analysed genomes. The maximum likelihood phylogenetic tree was integrated alongside with the presence and absence of genes. Each row corresponds to an isolate and each column corresponds to the variation in genes. The presence of genes is showed in the decreasing frequency. Empty space indicates the absence of genes.

A significant proportion of MGEs is derived from ST772 *S. aureus* accessory genome, that includes SCC*mec* element, prophages, pathogenicity island, plasmids, transposons, and insertion sequence (Fig 8). Overall, in ST772 *S. aureus* genome, MGEs are diverse. Composition of accessory genome analysis revealed that the composition accounts for a total of 2727 CDS (ranging between, 435-2613) in ST772 *S. aureus* genome. Variation in the accessory genome within subclade IIA, particularly results from the loss and gain of SCCmec V elements. A putative pathogenicity island carrying enterotoxin gene *sec* and *sel* was identified in all the genomes.

**Fig 8:**
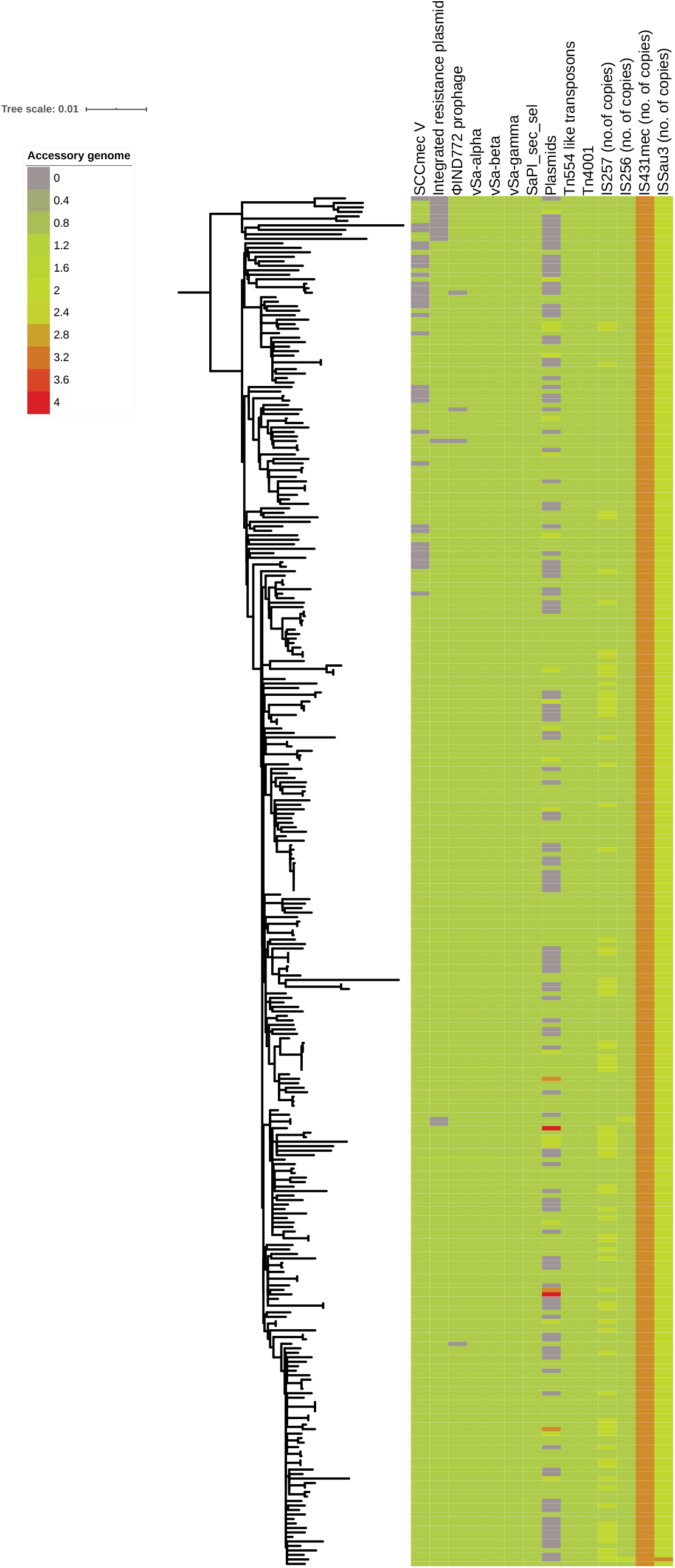
Mobile genetic elements identified in the accessory genome of ST772 *S. aureus* isolates.

## Discussion

The epidemiology of MRSA is changing and evolving constantly.^63^ Several well-characterised MRSA strains are predominant in different regions across the world. The present study revealed the genetic diversity of MRSA dominated by the internationally recognised CCs (CC1, CC22, and CC8). This highlights the risk of MDR and the significant threat of dissemination. In India, PVL positive ST772-SCC*mec* V and ST22-SCC*mec* IV are reported as the first and second most dominant lineages in causing MRSA infections.^16-18^ This was confirmed in the present study. These lineages have been reported from community and hospital settings. ^16-18,20,21,64-66^ The ST239-SCC*mec* III lineage has been reported as the predominant lineage in many Asian countries.^67^ In the present study, ST239-SCC*mec* III was found as a third most dominant MRSA lineage. Dissemination of ST772 and ST22 MRSA is gradually replacing the ST239-SCC*mec* III lineage in Indian hospitals. ^68^

In this study, a high degree of variability was seen among the PVL positive ST22 MRSA affects not only the SCC*mec* marker but also the core genome. The presence of unusual SCC*mec* types in ST22 MRSA indicates an evolutionary genetic change and so demands deeper genomic investigation. Studies have demonstrated the existence of multiple SCC*mec* types in MRSA.^69-71^ In the present study, 6% of the isolates had multiple SCC*mec* types. This suggests that multiple transfer events of SCC*mec* types would resulted in the integration of multiple *ccr* gene complexes in *S. aureus*. ^33, 72^

ST772-SCC*mec* V is outcompeting other *S. aureus* lineages in both community and hospital settings in India. Here, we provide evidence for the long term persistence of all subclades (IIA,B,C) of ST772 *S. aureus* and continued causing infections as reported earlier.^39^ This was evidence by the fact that each of the subclades contained atleast one isolate collected since early 2000s. Previously, three subclades (ST772-A1, A2, A3) were by defined by Steinig *et al*.^39^ Of the subclades, the subclade ST772-A1 is reported as old early branching lineage, whilst ST772-A2 and ST772-A3 are defined as dominant and more recent emerging subclades respectively. In this study, Indian ST772 *S. aureus* isolates (including study isolates) were distributed across the clades I and II.

The presence of multiple SCC*mec* types in the basal strains and isolates of subclade IIA indicates stable core genome, but recombination in SCC*mec* elements can occur, as reported earlier.^41^ This reassures that SCC*mec* element play a key role in adaptation and rendering the organism fitness to survive. Although, ST772 *S. aureus* has spread internationally, its expansion in India has not been previously studied. Our study provides evidence for the dominance of this lineage in India. In addition, isolates from other countries clustered closely with the Indian isolates, which indicate travel related spread.

The core genome of ST772 S. aureus remains stable suggesting the strength of these genomes in adapting to evolutionary pressures for persistence. The substitution rate of ST772 *S. aureus* (1.16 × 10^−6^) estimated in this study, is similar to the previous finding which reported 1.18 × 10^−6^ substitutions/site/year.^39^ The substitution rate of ST772 *S. aureus* lineage is much lower than the globally dominant *S. aureus* lineages ST22 (1.3 × 10^−6^), ST239 (1.6 × 10^−6^) and USA 300 clone (1.3 × 10^−6^).^23,73,74^ ST772 *S. aureus* lineage has relatively stable core genome and for evidence of microevolution. Our data shows temporal introduction of IRP with the sequential acqsuition of SCCmec V (5C2) and accumulation of double serine mutation in 1987 leading to the expansion of multi-drug resistant clones in 1987. However, in the previous study, the ST772 *S. aureus* has been reported to be emerged on the Indian subcontinent in the early 1960s and an emergence of dominant clade and its subgroups has reported in the early 1990s.^39^ In the present study, accuracy of MRCA prediction is improved using precise Bayesian model suggested by Bayes factor. Our study indicates that the earliest isolate (2004) was from South Asia (India and Bangladesh), which supports the earlier hypothesis suggesting that South Asia was the likely origin of this lineage.^39^ Acqsuition of IRP, a variant of SCC*mec* V (5C2) and the presence of double serine mutation (grlA-S80Y, gyrA-S84L) in the dominant clade II isolates were consistent with previous findings and could be the reason behind increased fitness and global expansion of this lineage. ^39^

Studies have emphasized the contribution of mobile genetic elements (MGEs) in the emergence of ST772-SCC*mec* V as a multi-drug resistant and highly virulent lineage. ^20,21,41^ The reference genome DAR4145, a representative of ST772 contains SCC*mec* V (5C2), PVL carrying ΦIND772 prophage, pathogenicity island (vSa-alpha, beta, and gamma) and an IRP. In the present study, comparative genomic analysis showed that these genetic elements are the conserved molecular markers of ST772 lineage, with few exceptions. Furthermore, comparative analysis of accessory genome showed that MGEs contributed for significant proportion of ST772 *S. aureus* genome. This observation reflects the contribution of MGEs in evolution and dissemination as described in ST22-SCC*mec* IV lineage^23^ and USA clones (300, 500, 500 like).^75^

A landmark study on the genomic investigation of the ST772 *S. aureus* detailed the key factors involved in its expansion, evolution and intercontinental transmission. This includes acquisition of IRP, fixation of a smaller SCC*mec* V (5C2) and replacement of mutation in *grl*A (S80F to S80Y).^39^ A similar findings was confirmed in the present study. An extrachromosomal plasmid in USA300 and an integrated element in the SCC*mec* of ST80-SCC*mec* IV lineage shared a similar genetic backbone of the IRP of ST722-SCC*mec* V lineage.^39^ The substitution S84L with S80Y is reported as the most common double serine mutation in *S. aureus*.^76^ Acquisition of PVL carrying ΦIND772 prophage and the pathogenicity islands has occurred even before the emergence of ST772-SCC*mec* V lineage. No significant difference in the cellular toxicity or biofilm formation have been reported between the basal strains and a single globally disseminated clade (ST772-A).^39^ Multi-drug resistance could have provided a strong fitness advantage for the ST772 lineage in hospital settings.

Multiple intercontinental spread from India to various countries in Europe, Australia and Asia have been reported. ^39^ In addition, outbreaks in the NICU in Ireland and a hospital in Norway have also documented. ^77,78^ However, intense endemic dissemination of this clone is not evident following intercontinental spread. ST772-SCC*mec* V has a similar pattern of dissemination as with that of other CA-MRSA such as USA300 clone and the ST80-SCCmec IV lineage.

In conclusion, ST772, ST22 and ST239 were observed as the predominant STs. ST772-SCC*mec* V was seen as the well-established lineage. Whilst, the occurrence of ST22 with various SCC*mec* types suggests an evolutionary genetic change and so demands deeper genomic investigation. ST772-SCC*mec* V (Bengal bay clone) was emerged on the Indian subcontinent in 1959 and diverged into a dominant clade in 1987. The acquisition of IRP, SCC*mec* V (5C2) and the double serine mutations associated with fluroquinolone resistance are likely to be the important attributes to the success of the ST772 lineage. The clinical antimicrobial use pattern may have driven the spread and survival of ST772 MRSA in hospitals. ST772-SCC*mec* has the multi-drug resistance trait of HA-MRSA and the epidemiological characteristics of CA-MRSA.

## Acknowledgements

None

## Transparency declarations

None to declare

---

**Fig S1:**
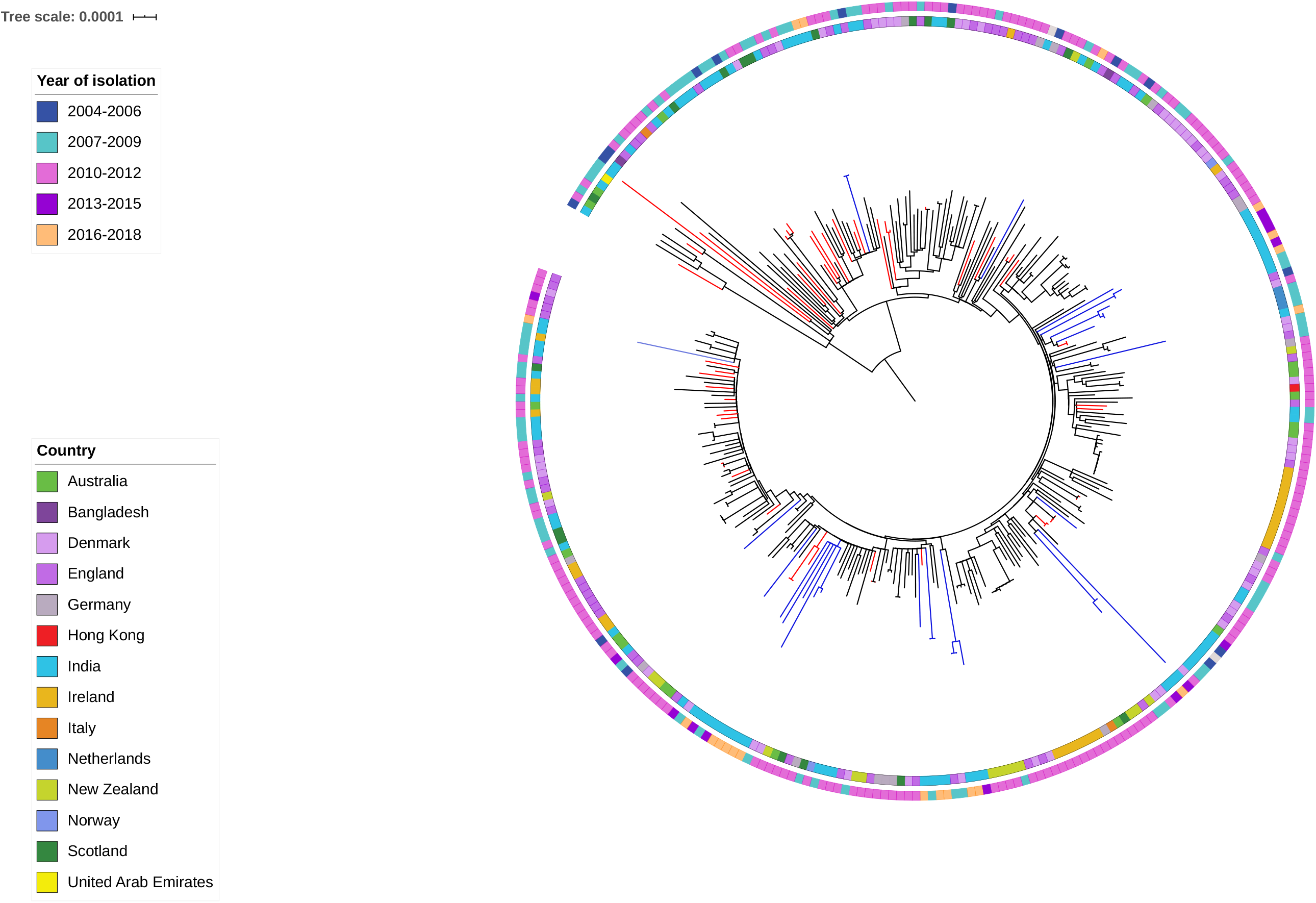
Maximum likelihood phylogenetic tree was constructed based on the core genome SNP of ST772 *S. aureus* lineage (n=304). The year and country of isolation of ST772 S. aureus isolates.

**Fig S2:**
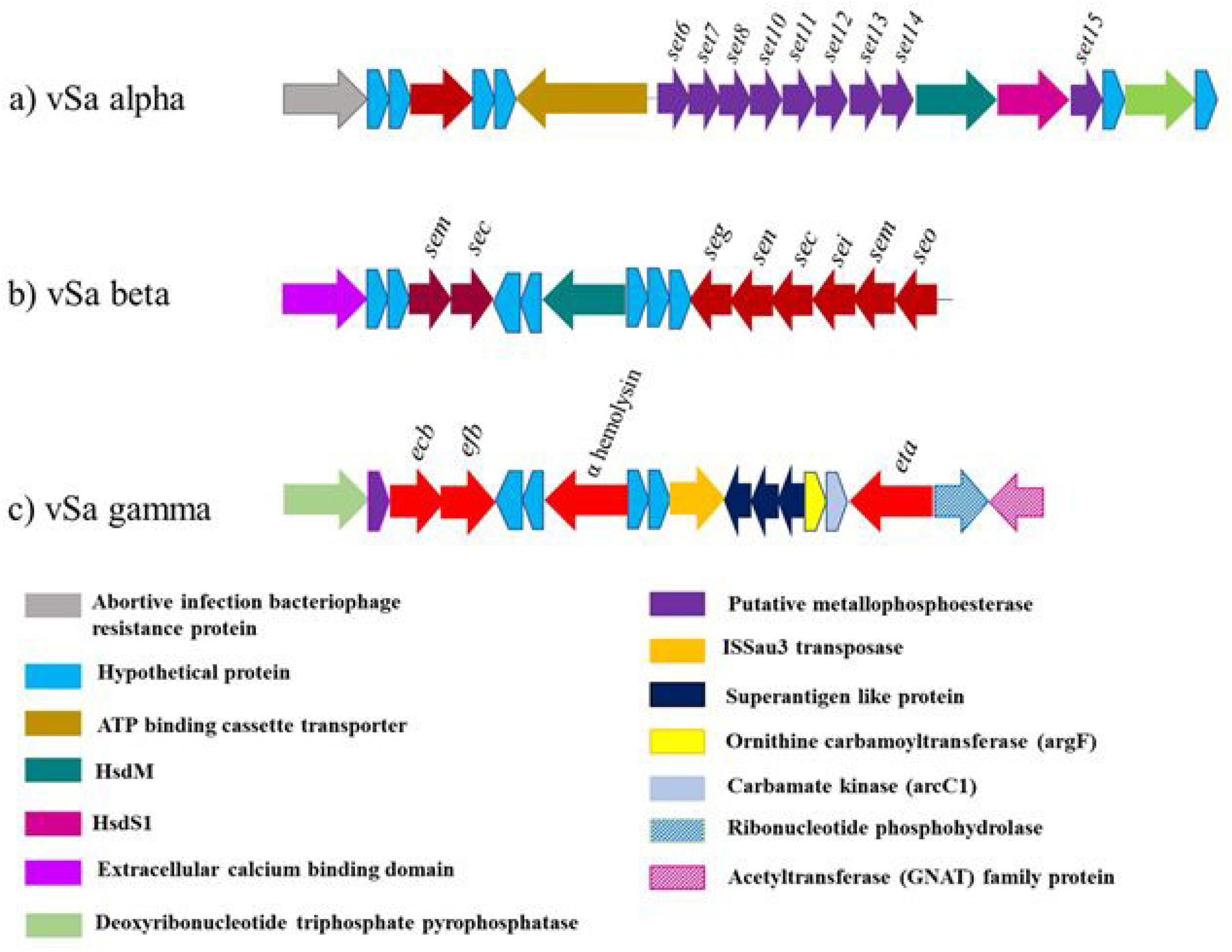
Pathogenicity island identified in ST772-SCCmec V lineage. a) vSa-alpha pathogenicity island carried various exotoxins and superantigens. b) vSa-beta pathogenicity island harboured enterotoxin gene cluster. c) vSa-gamma genomic island comprised of alpha hemolysin and exfoliative (*eta*) toxin.

**Fig S3:**
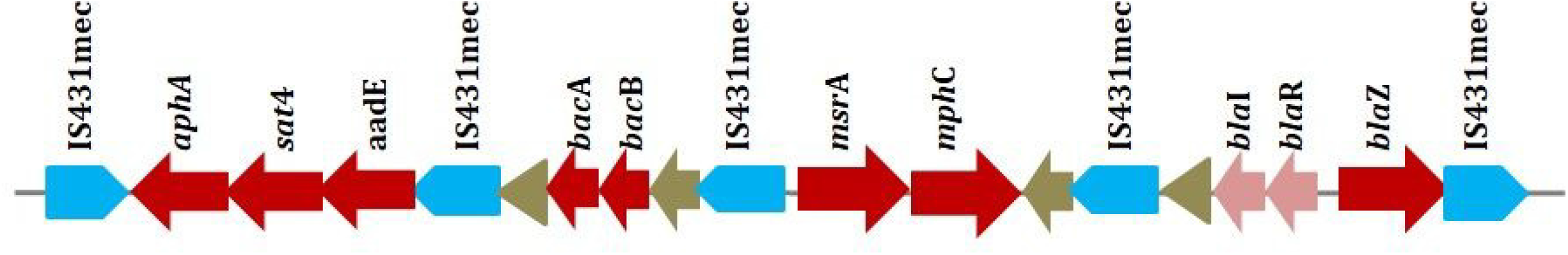
Genetic environment of integrated resistance plasmid (IRP) in VB9352 (accession no. CP035670)

